# Evaluation of 3D markerless motion capture accuracy using OpenPose with multiple video cameras

**DOI:** 10.1101/842492

**Authors:** Nobuyasu Nakano, Tetsuro Sakura, Kazuhiro Ueda, Leon Omura, Arata Kimura, Yoichi Iino, Senshi Fukashiro, Shinsuke Yoshioka

## Abstract

There is a need within human movement sciences for a markerless motion capture system, which is easy to use and sufficiently accurate to evaluate motor performance. This study aims to develop a 3D markerless motion capture technique, using OpenPose with multiple synchronized video cameras, and examine its accuracy in comparison with optical marker-based motion capture. Participants performed three motor tasks (walking, countermovement jumping, and ball throwing), with these movements measured using both marker-based optical motion capture and OpenPose-based markerless motion capture. The differences in corresponding joint positions, estimated from the two different methods throughout the analysis, were presented as a mean absolute error (MAE). The results demonstrated that, qualitatively, 3D pose estimation using markerless motion capture could correctly reproduce the movements of participants. Quantitatively, of all the mean absolute errors calculated, approximately 47% were less than 20 mm and 80% were less than 30 mm. However, 10% were greater than 40 mm. The primary reason for mean absolute errors exceeding 40mm was that OpenPose failed to track the participant’s pose in 2D images owing to failures, such as recognition of an object as a human body segment, or replacing one segment with another depending on the image of each frame. In conclusion, this study demonstrates that, if an algorithm that corrects all apparently wrong tracking can be incorporated into the system, OpenPose-based markerless motion capture can be used for human movement science with an accuracy of 30mm or less.

## 1. Introduction

Motion capture systems have been used extensively as a fundamental technology within biomechanics research. However, traditional marker-based approaches experience significant environmental constraints. For example, measurements are diffcult to perform in environments wherein wearing markers during the activity is not ideal (such as sporting games). Markerless measurements without such environmental constraints can facilitate new learnings about human movements (Mündermann et al., 2006); however, complex information processing technology is required to make an algorithm that recognizes human poses or skeletons from images. Therefore, it is desirable to many biomechanics researchers to develop a markerless motion capture that is easy to use.

Recently, automatic human pose estimation using deep learning techniques have attracted attention amongst computer vision researchers. OpenPose is one of the most popular technologies (Cao et al., 2018) and is deemed easy to use for biomechanics researchers. It is open source software that automatically estimates human joint centers and skeletons from 2D RGB images, outputting the 2D coordinates in the images. Kinect is another easy-to-use markerless motion capture that has been used in many studies (Clark et al., 2012; Pfister et al., 2014; Gao et al., 2015; Schmitz et al., 2014). OpenPose, when compared to Kinect, has less constraints on the distance between the camera and the target to be measured as well as the sampling rate of video recording, because it can estimate the human pose from RGB image without using a depth sensor.

Seethapathi et al. (2019), which reviewed pose tracking studies from the perspective of movement science, pointed out that human pose tracking algorithms, such as OpenPose, did not prioritize the quantities that matter for movement science. It remains unclear whether the accuracy of the OpenPose-based 3D markerless motion capture is appropriate for human movement studies such as sports biomechanics or clinical biomechanics. The aim of this study is to develop a 3D markerless motion capture using OpenPose with multiple synchronized video cameras, then assess the accuracy of the 3D markerless motion capture by comparing with an optical marker-based motion capture.

## 2. Materials and Methods

### Participants

Two healthy male volunteers participated in this experiment. The mean age, height, and body mass of the participants were 22.0 years, 173.5 cm, and 69.5 kg, respectively. The participants provided written informed consent prior to the commencement of the study, and the experimental procedure used in this study was approved by the Ethics Committee of the university with which the authors were affliated.

### Overview of data collection

Participants performed three motor tasks in the following order: walking, countermovement jumping, and ball throwing. These movements were measured using both a marker-based optical motion capture and a video camera-based (markerless) motion capture. A light was used to synchronize the data obtained from all the video cameras and the two different measurement systems. All methods and instrumentation details are in the following subsections.

### Marker-based motion capture

Forty-eight reflective markers were attached onto body landmarks (Figure 1). The coordinates of these reflective markers upon the participants’ bodies were recorded using a 16-camera motion capture system (Motion Analysis Corp, Santa Rosa, CA, USA) at a sampling rate of 200 Hz. The elbow, wrist, knee, and ankle joint centers were assigned to the mid-points of the lateral and medial markers, while the shoulder joint centers were assigned to the mid-points of the anterior and posterior shoulder markers. The hip joint centers were estimated using the method described by Harrington et al. (2007). The raw kinematic data was smoothed using a zero-lag fourth order Butterworth low-pass filter. The cut-off frequency of the filter was determined using a residual analysis (Winter, 2009). Data analysis was performed using MATLAB (v2019a, MathWorks, Inc., Natick, MA, USA).

**Figure 1:**
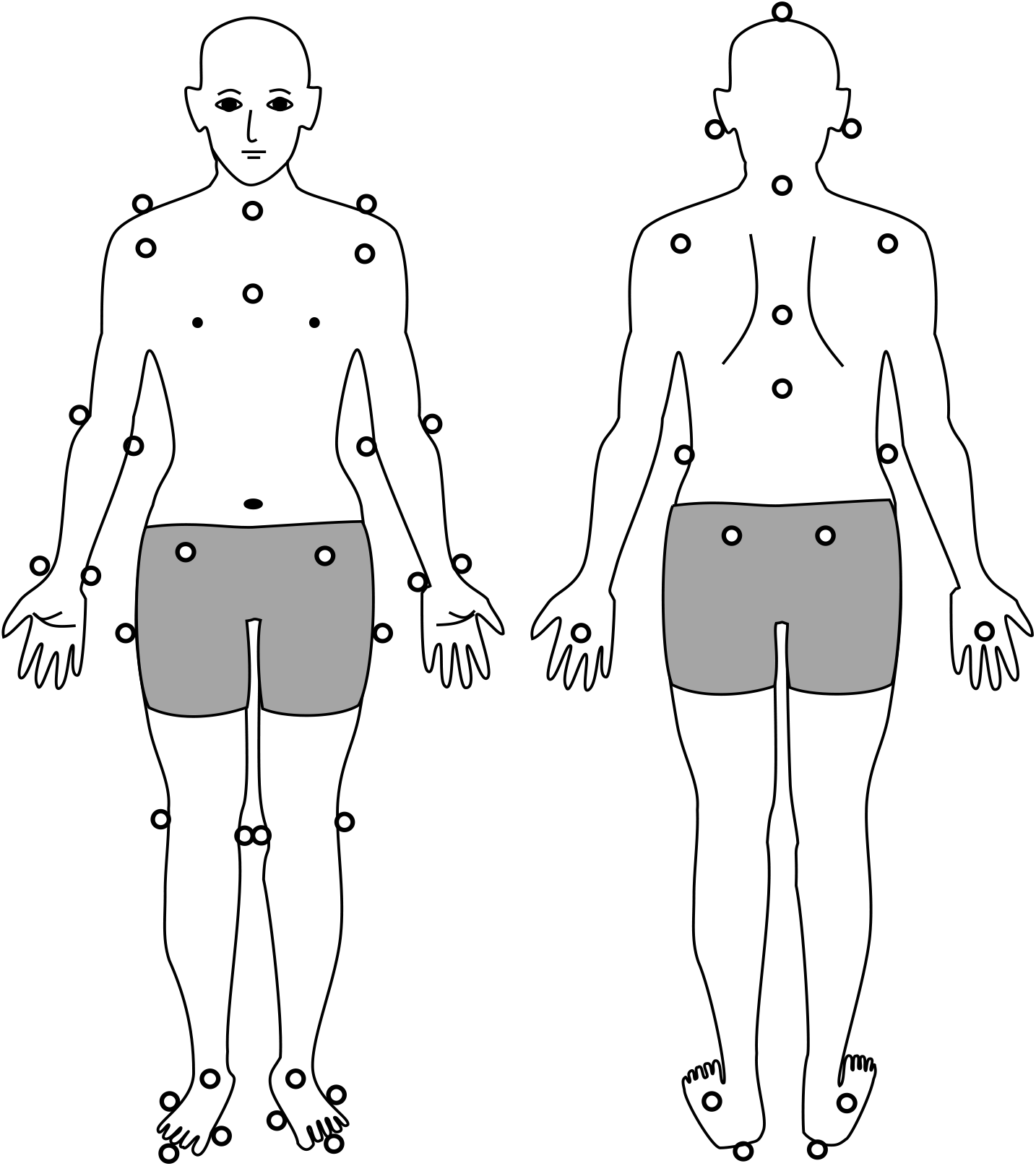
Positions of reflective markers attached on the landmarks of the body. Forty-eight markers were placed on the fingertip (right), third metacarpal, styloid process of ulna, styloid process of radius, humerus-medial epicondyle, humerus-lateal epicondyle, humerus-lesser tubercle, under the scapula-acromial angle, scapula-acromion, toe, first metatarsal, fifth metatarsal, calcaneus, malleolus medialis, malleolus lateralis, knee medial side, knee lateral side, trochanter major, top of head, ear, upper margin of sternum, C7 vertebra, lowest edge of rib, sternum-xiphoid process, T8/12 vertebra, anterior superior iliac spine and posterior superior iliac spine.

### Markerless motion capture

The experimental setup and overview of the markerless motion capture are shown in Figure 2. The markerless motion capture consisted of five video cameras (GZ-RY980, JVCKENWOOD Corp, Yokohama, Kanagawa, Japan). Two measurement conditions, *i.e.* combinations of video camera resolutions and sampling frequencies, were implemented: 1920 × 1080 pixels at 120 Hz (1K condition) and 3840 × 2160 pixels at 30 Hz (4K condition). OpenPose (version 1.4.0) was installed from GitHub (CMU-Perceptual-Computing-Lab, 2017) and run with GPU (GEFORCE RTX 2080 Ti, Nvidia Corp, Santa Clara, CA, USA) under default settings. Twenty-five keypoints (Figure 3) of the participant’s body were outputted independently for each frame. The control points, at which 3D global coordinates could be identified, were measured using the video cameras with use of a calibration pole. The 2D video camera coordinates obtained from OpenPose were transformed to 3D global coordinates using a direct linear transformation (DLT) method (Miller et al., 1980). The raw kinematic data was smoothed using a zero-lag fourth order Butterworth low-pass filter. The cut-off frequency of the filter was determined using residual analysis (Winter, 2009) with its ranges of 5-8 Hz and 2-3 Hz in the 1K and 4K conditions, respectively.

**Figure 2:**
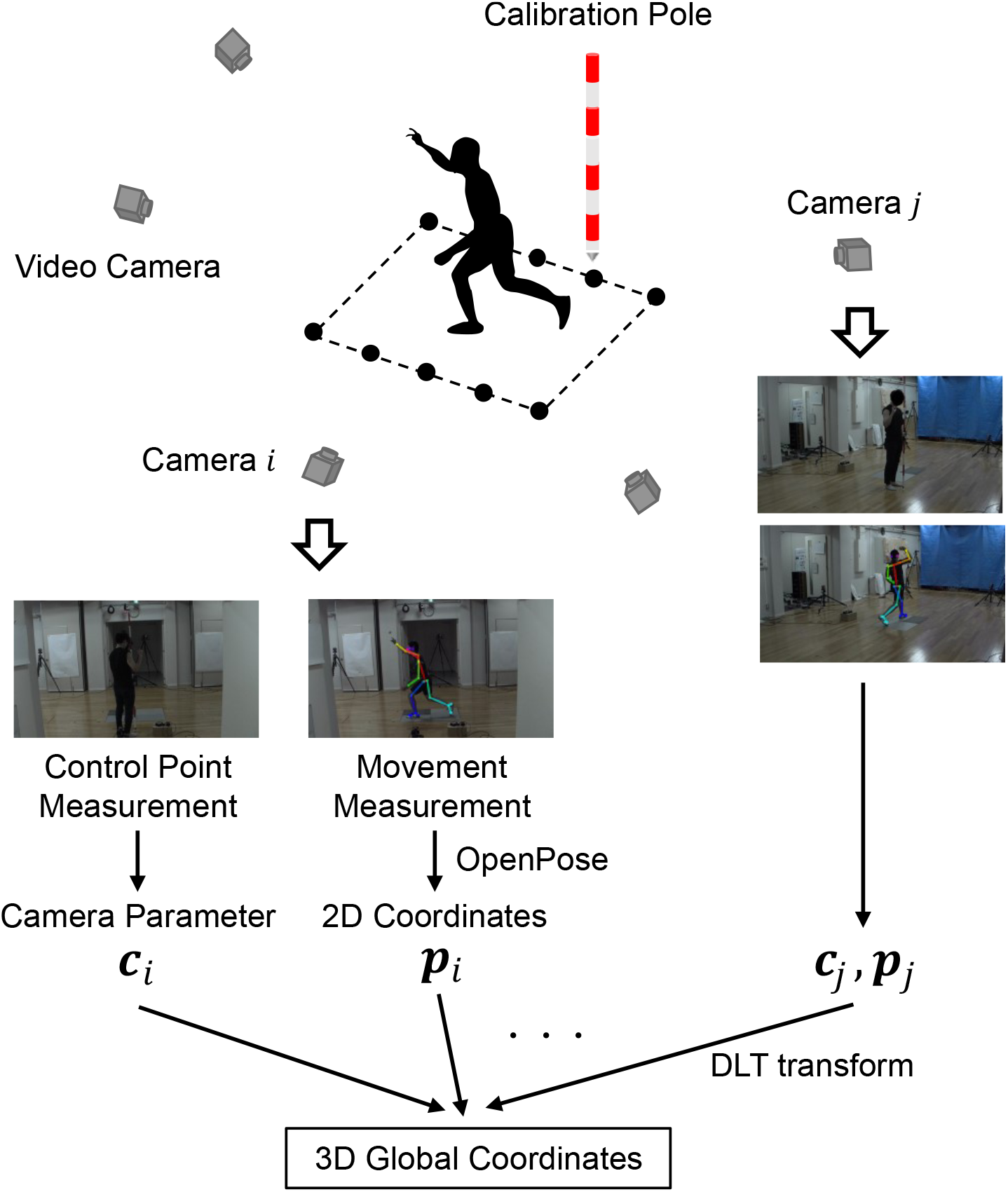
Experimental setup and overview of the markerless motion capture.

**Figure 3:**
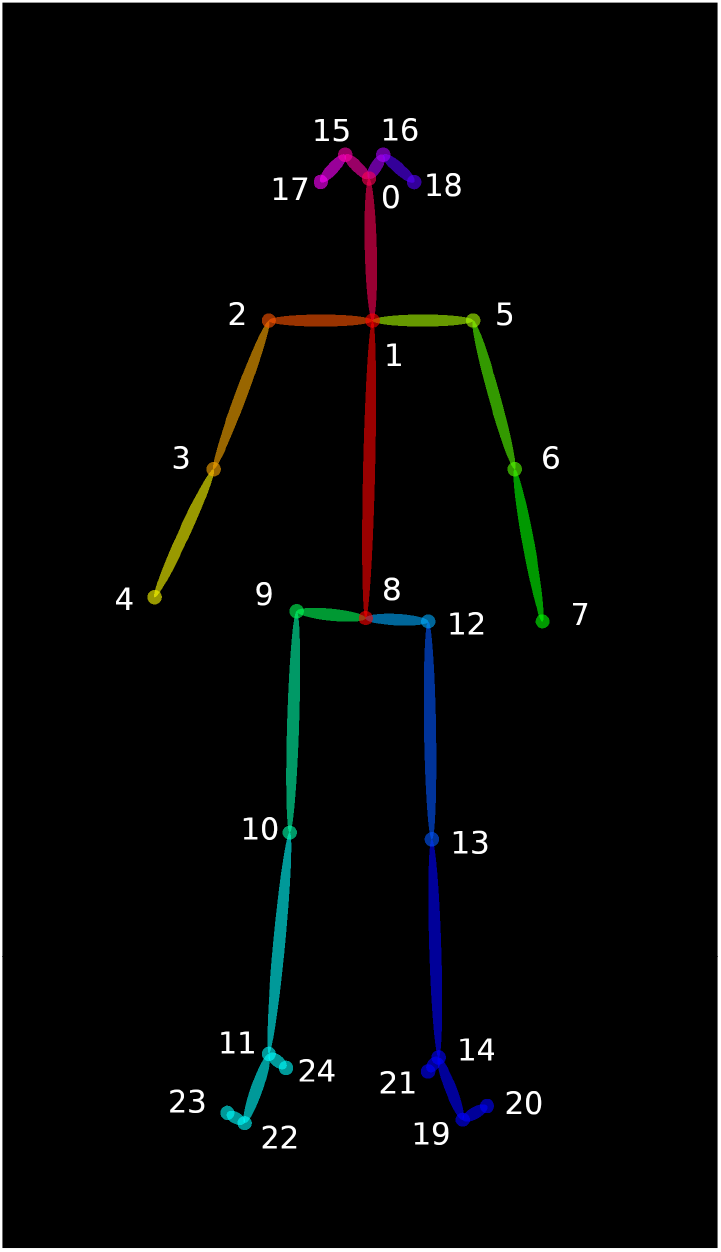
Twenty-five keypoints (nose, neck, shoulder, elbow, wrist, hip, knee, ankle, eye, ear, big-toe, small-toe, and heel) of human body tracked by OpenPose (CMU-Perceptual-Computing-Lab, 2017).

### Data analysis

The position data obtained using the marker-based motion capture was downsampled using the spline function to alter the number of frames such that they are the same as that obtained using markerless motion capture. The analysis period durations were defined for each individual motor task as follows: from the second step heel contact to the next heel contact of the same leg in a walking task, from the start of the squatting motion to the recovery of the initial upright stance in a jumping task, and from the toe-off on the opposite side of the throwing arm to the end of the arm-swing in a throwing task. The differences in the corresponding joint positions that were estimated from the two different motion captures throughout the analysis durations were calculated. Mean absolute error (MAE) was used as the indicator of the difference as described by Equation 1, where, *x*_*m*_ and *x*_*o*_ are the positions estimated by the marker-based and OpenPose-based approaches, respectively.

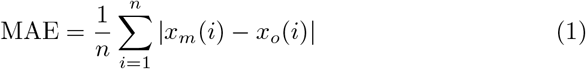

## 3. Results

Examples of the 3D pose estimations obtained by the two different motion captures are depicted in Figure 4. In addition, video examples that show the participant’s pose during movements are provided as supplementary materials to this paper. The landmarks’ positions tracked by OpenPose (Figure 3) do not necessarily correspond to the points estimated by the marker-based approach. The joint positions of the shoulder, elbow, wrist, hip, knee, and ankle tracked by OpenPose are considered to be approximately the same as those estimated by the marker-based approach. However, the positions of other landmarks tracked by OpenPose are considered to be different from those estimated by the marker-based approach. The representative time-series profiles of joint positions estimated by both the marker-based motion capture (Mocap) and the OpenPose-based markerless motion capture (OpenPose) can be seen in Figure 5. The mean absolute error (MAE) of the two plots throughout the duration of analysis is shown in each panel. The shapes of the time-series profiles were found to be approximately the same.

**Figure 4:**
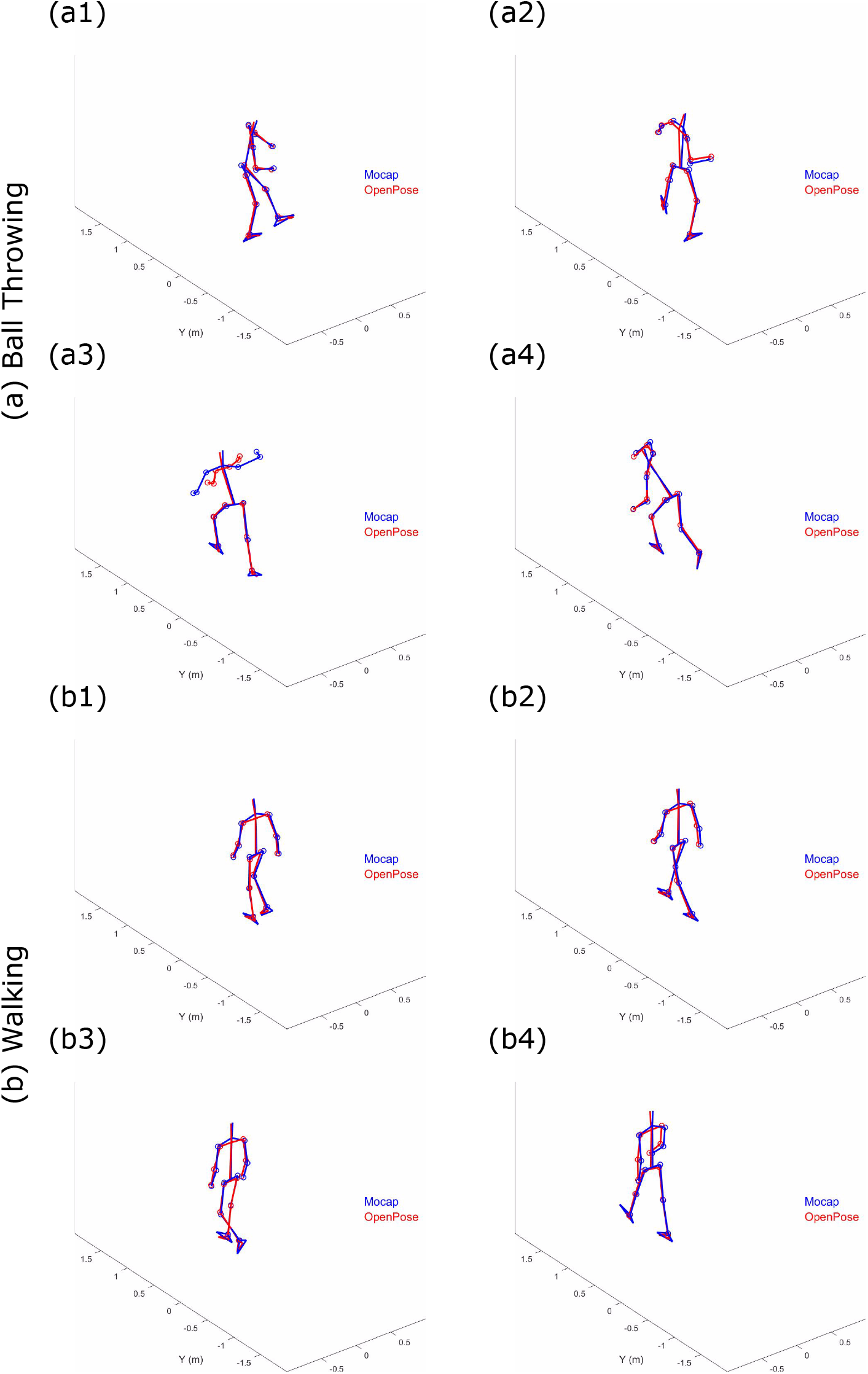
Examples of participant’s pose estimated by the marker-based motion capture (Mocap) and by the markerless motion capture using OpenPose (OpenPose) during a (a) ball throwing task and (b) walking task.

**Figure 5:**
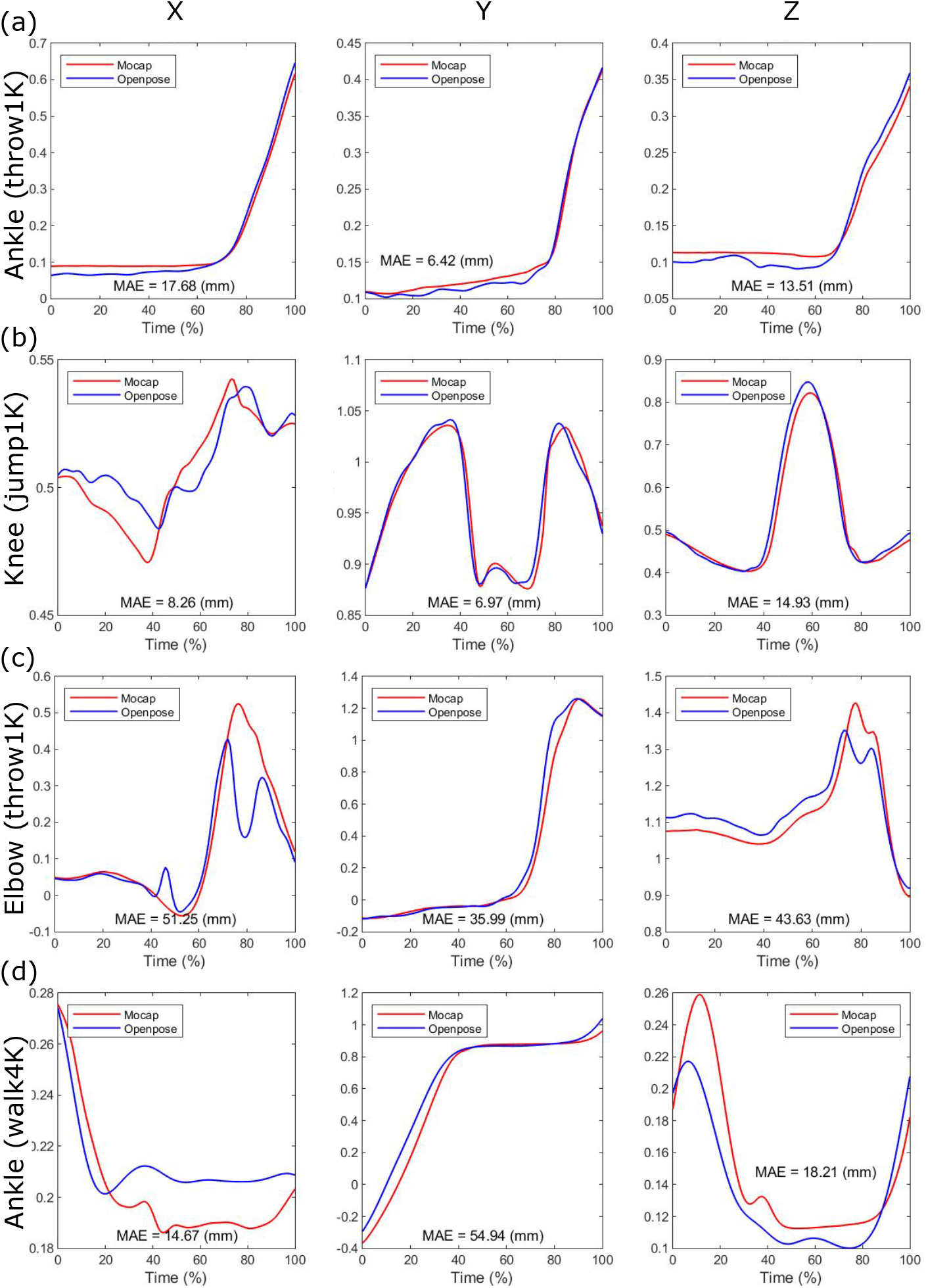
Time series profiles of joint positions estimated by the marker-based motion capture (Mocap) and by the markerless motion capture using OpenPose (OpenPose). The mean absolute error (MAE) through analysis duration is shown in each panel.

Quantitatively, the MAE of joint positions in Figure 5a and Figure 5b were less than 20 mm; however, the MAEs of joint positions in Figure 5c and Figure 5d were greater than 40 mm. The MAEs were particularly large at specific moments within the analysis duration (*i.e.* at 45% and 80% time in Figure 5c, described in Figure 4c). The MAEs of the corresponding joint positions, estimated from the two different motion captures for all trials, are presented in Table 1. Of these MAEs in Table 1, approximately 47% are less than 20 mm, 80% are less than 30 mm, and 10% is greater than 40 mm.

**Table 1:** The differences of corresponding joint positions estimated from the two different motion captures.

|  |  | Walk<sub>1K</sub> | Walk<sub>4K</sub> | Jump<sub>1K</sub> | Jump<sub>4K</sub> | Throw<sub>1K</sub> | Throw<sub>4K</sub> |
| --- | --- | --- | --- | --- | --- | --- | --- |
| ShoulderR | X | 28.4 | 24.6 | 27.1 | 20.9 | 21.7 | 19.3 |
|  | Y | 17.0 | 49.7 | 12.5 | 11.9 | 27.7 | 32.5 |
|  | Z | 17.9 | 16.5 | 29.5 | 30.8 | 15.8 | 13.9 |
| ElbowR | X | 4.32 | 6.96 | 9.36 | 8.25 | 47.3 | 45.2 |
|  | Y | 37.0 | 66.8 | 27.0 | 20.5 | 28.7 | 35.1 |
|  | Z | 21.7 | 22.2 | 26.9 | 35.3 | 38.0 | 38.9 |
| WristR | X | 5.78 | 7.52 | 8.96 | 8.31 | 40.6 | 47.5 |
|  | Y | 19.0 | 44.2 | 13.2 | 20.6 | 28.7 | 40.3 |
|  | Z | 15.7 | 16.8 | 23.8 | 38.9 | 24.7 | 26.7 |
| HipR | X | 9.65 | 7.67 | 8.77 | 6.01 | 29.5 | 25.0 |
|  | Y | 21.3 | 49.4 | 15.0 | 14.2 | 13.5 | 20.8 |
|  | Z | 24.4 | 20.6 | 31.0 | 32.1 | 27.2 | 23.5 |
| KneeR | X | 6.41 | 4.09 | 7.74 | 6.47 | 15.2 | 13.1 |
|  | Y | 25.9 | 48.2 | 8.48 | 18.3 | 13.8 | 19.1 |
|  | Z | 10.1 | 11.4 | 14.8 | 20.9 | 24.4 | 20.4 |
| AnkleR | X | 9.68 | 8.73 | 9.82 | 6.67 | 12.3 | 19.1 |
|  | Y | 28.6 | 58.1 | 9.31 | 11.0 | 17.7 | 22.2 |
|  | Z | 11.7 | 20.7 | 20.6 | 27.9 | 12.4 | 20.3 |

The accuracy of the 3D pose estimation using the markerless motion capture depends on 2D pose tracking by OpenPose. Because the algorithm that tracks the human pose was applied to each frame of the video independently, within a single trial, there are frames where the participant’s pose was well tracked, whereas in others, the participant’s pose was not well tracked. Examples of pose estimation successes and failures using OpenPose are depicted in Figure 6.

**Figure 6:**
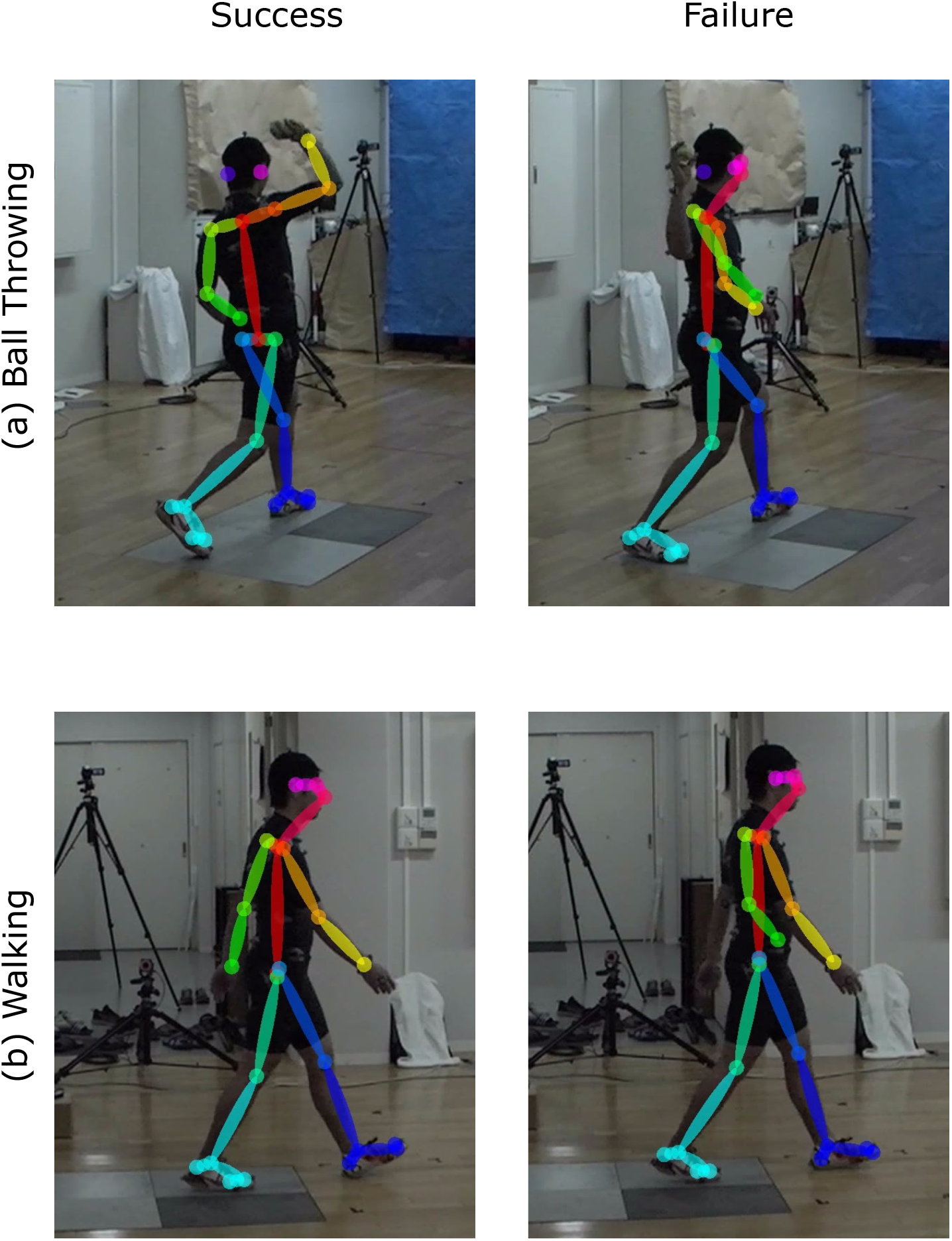
Examples of pose estimation using OpenPose during a (a) ball throwing task and (b) walking task. Left panels show the frame where the tracking of all joints was succeeded (to some extent), and right panels show the frame where the tracking of right arm joints was missed.

## 4. Discussion

This study aimed to examine the accuracy of 3D markerless motion capture using OpenPose with multiple video cameras, through comparison with an optical marker-based motion capture. Qualitatively, 3D pose estimation using the markerless motion capture approach can correctly reproduce the movements of participants (Figure 4 and Supplementary movies). The MAE in 80% of all trials conducted was found to be less than 30 mm. This small error may be due to the shortcomings in the OpenPose tracking precision. Because the majority of computer vision-based pose tracking algorithms, including OpenPose, are based on the supervised learning using manually labeled data, it is inevitable that small errors in 3D pose are caused by inherent noise in the training data.

Proportionately large MAEs exceeding 40 mm were observed in certain cases (Figure 5). Observing the estimated pose during movements reveals that when the correct joint positions (including noises) were estimated in the trial, an error of less that 30 mm existed (*e.g.* Figure 4 a1, a2, a4). However, when apparently incorrect joint positions were estimated, a comparatively large error of greater thatn 40 mm was observed (*e.g.* Figure 4 a3).

Observing the estimated pose during movements reveals that the correct joint positions (including noises) are estimated in the trial including a small error that is less than 30 mm (*e.g.* Figure 4 a1,a2,a4); however, apparently incorrect joint positions are estimated in the trial including a relatively large error that is more than 40 mm (*e.g.* Figure 4 a3). The primary reason for estimating apparently incorrect 3D positions is that OpenPose failed to track the participant’s pose depending on the image of each individual frame (Figure 6). Du to the 2D tracking failures, correction of such failures is required to achieve a more accurate 3D pose estimation. Within this study, for example, the interchange of the left and right segments was retrospectively corrected because without this correction, the 3D pose was completely different from the human shape. However, recognizing an object as a human body segment (*e.g.* failures in Figure 6) was not corrected because it may have required manual tracking; moreover, without the correction, the 3D pose accuracy could be evaluated. Therefore, to use the OpenPose-based markerless motion capture in human movement science studies, it is considered necessary to incorporate algorithms that can correct all such a tracking failures.

Other sources of error may be data processing, such as time synchronization. The error in the movement direction (*i.e.* Y direction for walking task and Z direction for jumping task), especially in the 30 Hz measurement (4K condition), tends to be large (Table 1). Within the time series profiles, the timing of synchronization appears to affect to the error of the two motion captures (Figure 5d). However, because this is a problem caused by the process of comparing the two motion captures, the effect of this error on the accuracy of the markerless system measurement should be relatively small.

This study is preliminary work, and thus requires further examination. The accuracy of other biomechanical parameters, such as joint angle, joint angular velocity, and joint torque, needs to be investigated. Additionally, a program that is capable of correcting the OpenPose tracking failures described in this study, which can improve the accuracy of the 3D estimation, needs to be developed. While limitations still exist, the OpenPose-based markerless motion capture is expected to be applied in the future to sporting games that have been considered diffcult to measure with marker-based motion capture.

## Acknowledgements

This work was supported by JST-Mirai Program Grant Number JPMJMI18C7, Japan.

## Supplementary material

Supplementary videos are attached with this paper. The videos show the human pose during walking, jumping, and throwing obtained by two different motion capture.

